## Supplementary for "Extracellular adenosine deamination primes tip organizer development in *Dictyostelium*"

### Supplementary Figure 1

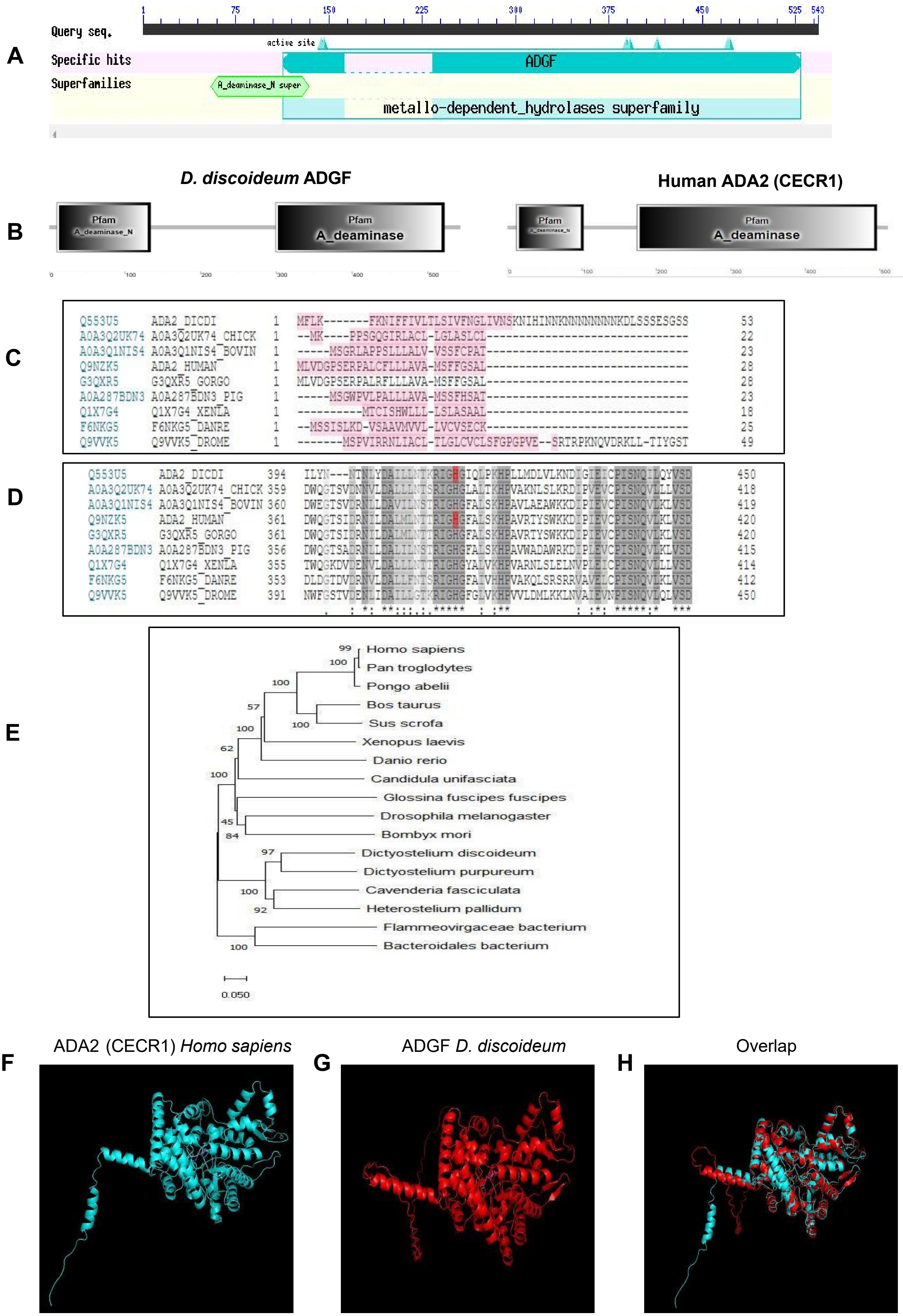

**Figure S1. Bioinformatic analyses of ADGF**

**A)** BLAST analysis of ADGF. **B)** SMART analysis depicting different ADA domains within ADGF. ADGF has an ADA and an N-terminal deaminase domain similar to human ADA2. **Multiple sequence alignment of *Dictyostelium* ADGF with ADGF from other organisms. C)** The shaded region depicts the N-terminal signal sequence characteristic of extracellular proteins. **D)** The active site residue highlighted in red, is conserved between *D.discoideum* and human *ada2*. **E)** Phylogenetic analysis of ADGF across different organisms. Maximum likelihood method was used for constructing the tree using molecular evolutionary genetic analysis X (MEGAX ). **Structural comparison of human ADA2 and *Dictyostelium* ADGF**. Identical tertiary structures of human ADA2 and *Dictyostelium* ADGF. **F)** ADA2 (CECR1) *Homo sapiens* and **G)** DdADGF **H)** Alignment of *Dictyostelium* ADGF with Human ADA2 (CECR1).

### Supplementary Figure 2

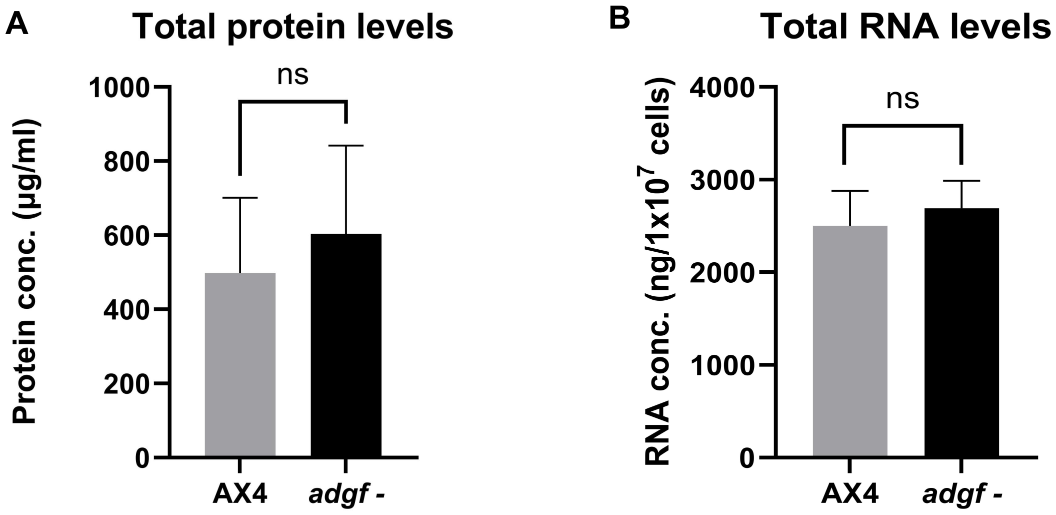


**Figure S2. Total protein and RNA levels during mound stage of development.**

**A)** Total protein levels were estimated using BCA assay. **B)** Total RNA levels in the WT and *adgf*⁻. The error bars represent the mean ± SEM (n=3). ns = not significant.

### Supplementary Figure 3

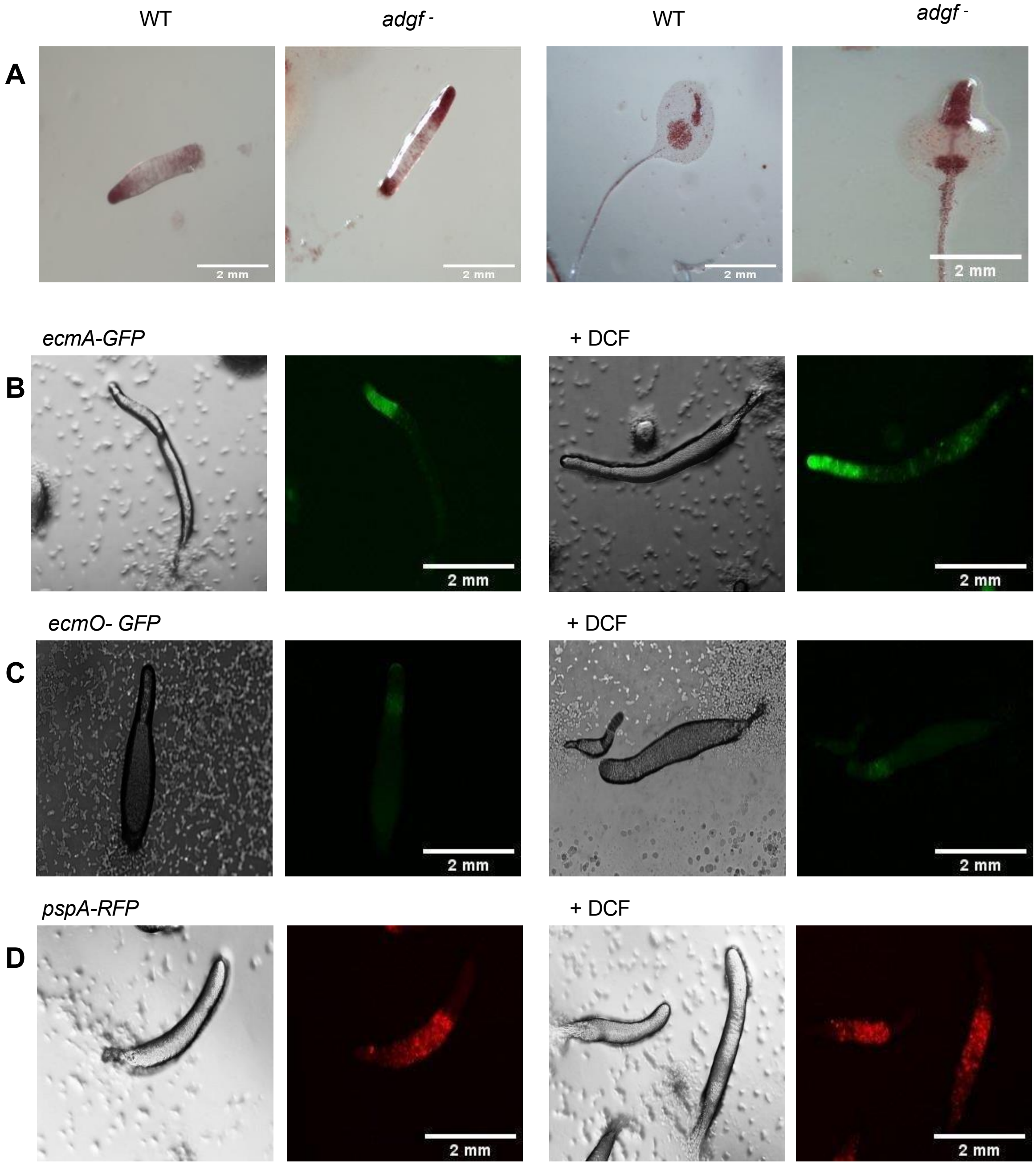


**Figure S3. Neutral red staining of mounds and slugs.**

**A)** WT and *adgf* - cells were stained with NR, plated and images were captured at different time intervals. Scale bar: 2 mm; (n=3). **Pst/psp marker expression in slugs after inhibitor treatment. B.** *ecmA*-GFP. **C)** *ecmO*-GFP. **D)** *pspA*-RFP expression in the slugs after DCF treatment. Scale bar: 2 mm.

### Supplementary Figure 4

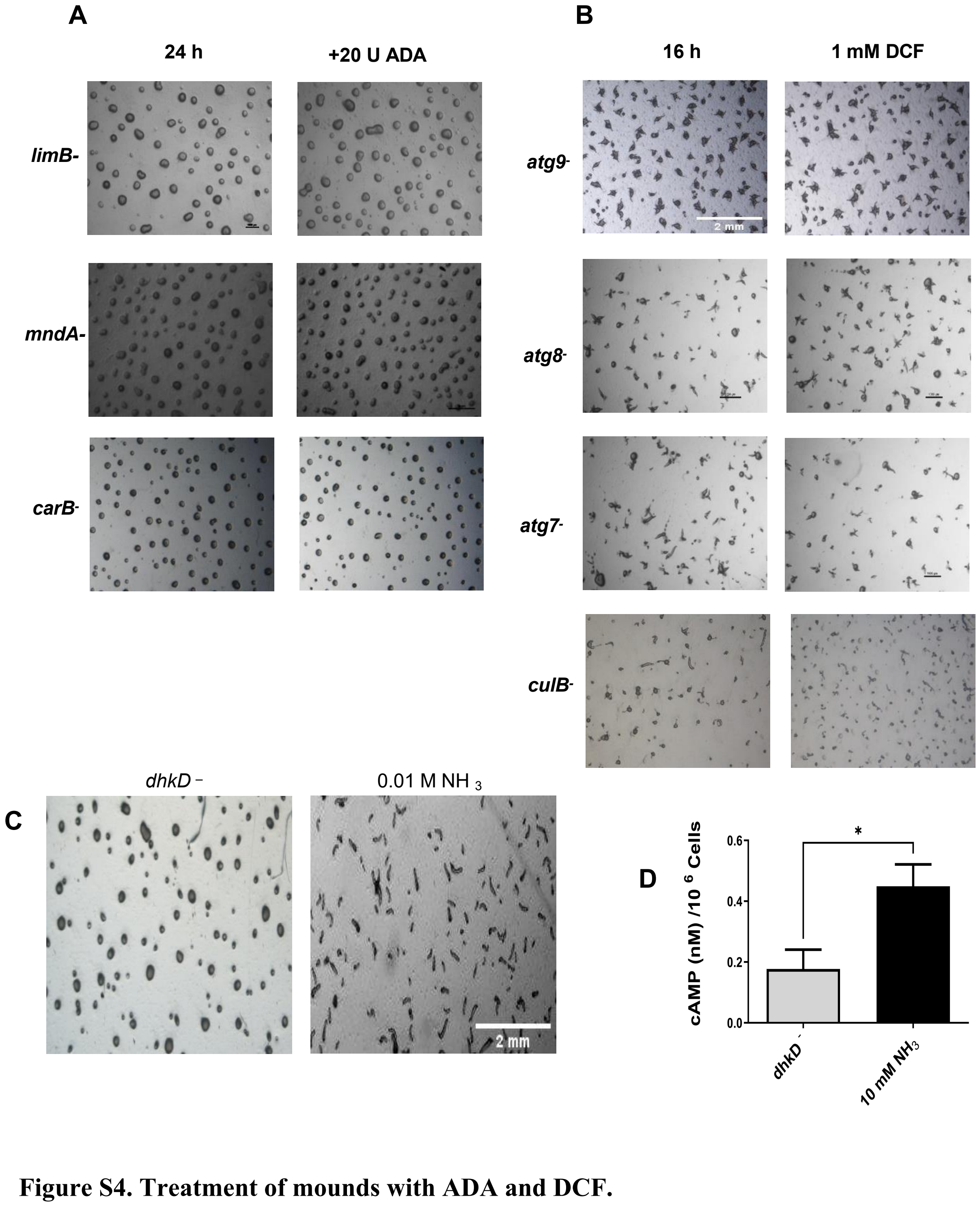


**Figure S4. Treatment of mounds with ADA and DCF.**

**A)** ADA enzyme was directly added on top of the mound arrest mutants at 12 h, and images were taken after 3.5 h. Scale bar: 2 mm; n=3. **B)** Multi tipped mutants were treated with 1 mM DCF, plated for development and images were taken after 16 h. Scale bar: 2 mm; n=3. **C)** Ammonia restores tip formation in *dhkD*$^{-}$ mounds.. Scale bar: 2 mm; n=3. **D)** cAMP levels in *dhkD* mutants rescued with ammonia.

### Supplementary Figure 5

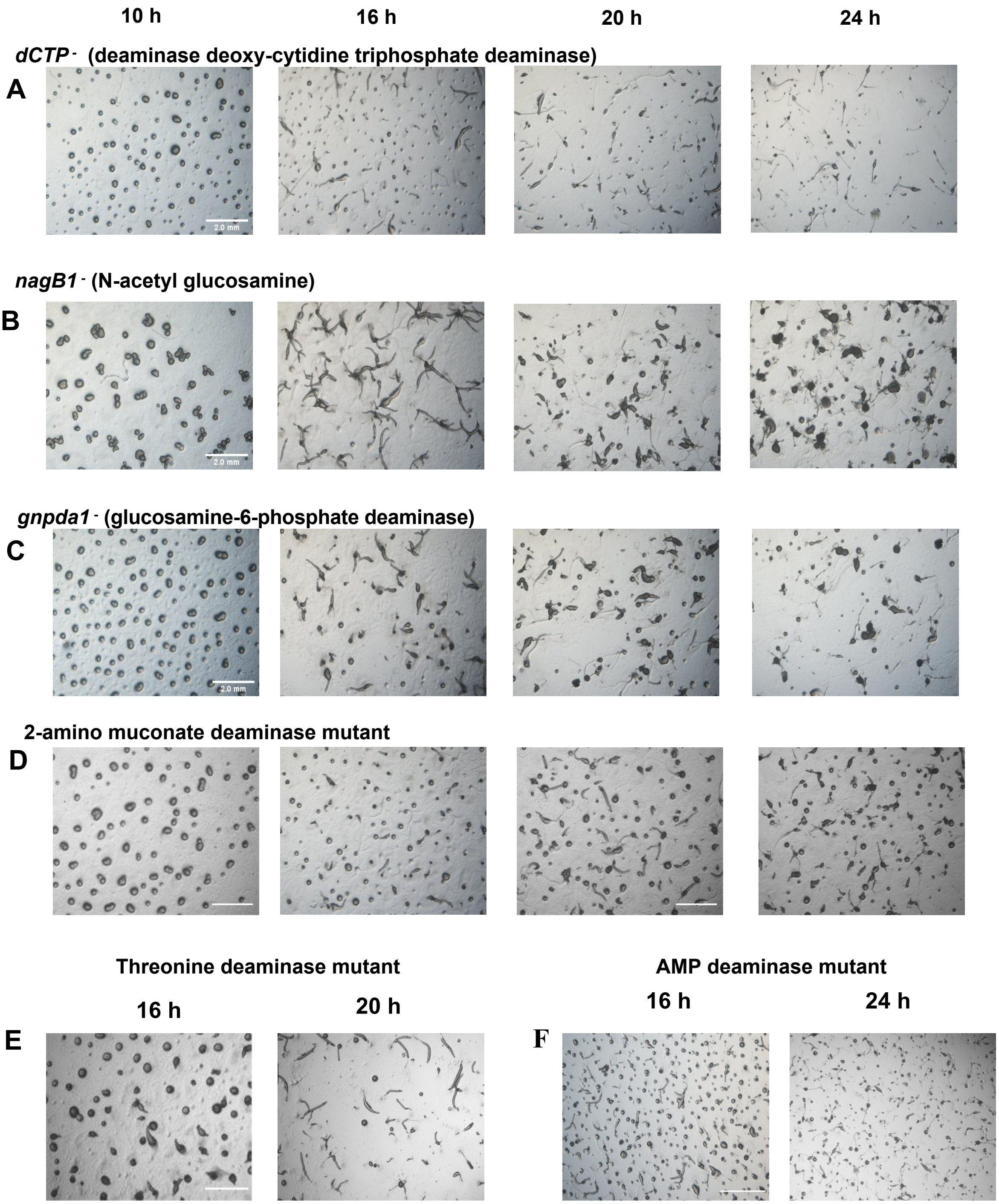


**Figure S5. Developmental phenotype of different deaminase gene knockouts .**

**A)** deoxy-cytidine triphosphate deaminase **B)** N-acetyl glucosamine **C)** glucosamine-6-phosphate deaminase **D)** 2-amino muconate deaminase. **E)** Threonine deaminase **F)** Adenosine monophosphate deaminase. Scale bar: 2 mm; n=3.

### Supplementary Figure 6

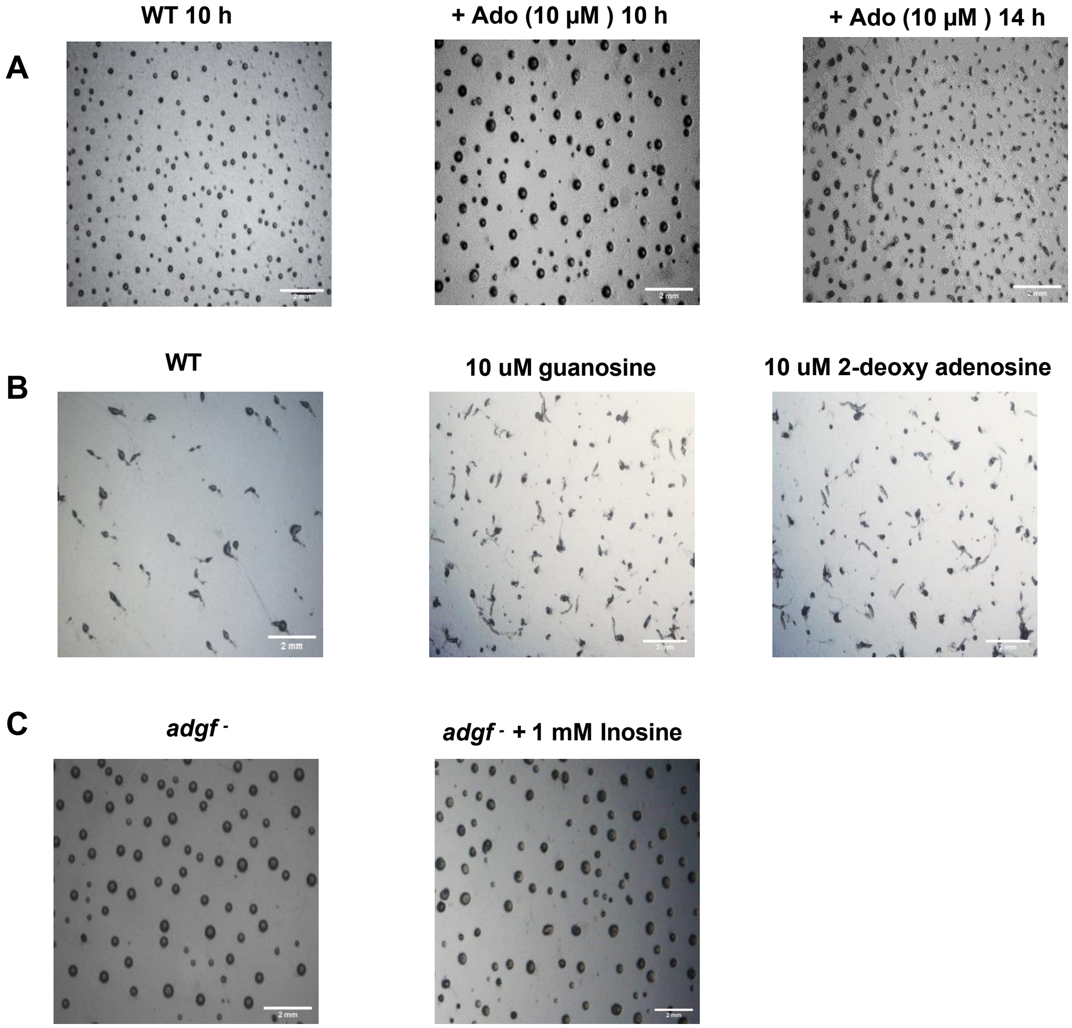


**Figure S6. Treatment with adenosine and other purines does not induce mound arrest in WT.**

**A)** WT cells were treated with adenosine (10 µM) while plating, and imaged at different time intervals. Scale bar: 1 mm; n=3. **B)** WT cells were treated with 10 µM guanosine and 10 µM 2-deoxy adenosine, plated for development and images were taken after 16 h. Scale bar: 2 mm; n=3. **C)** *adgf* $^-$ mounds were treated with inosine and images were captured 3.5 h post chemical treatment. Scale bar: 2 mm; (n=3).

### Supplementary Figure 7

*adgf* - developed in the presence of bacteria

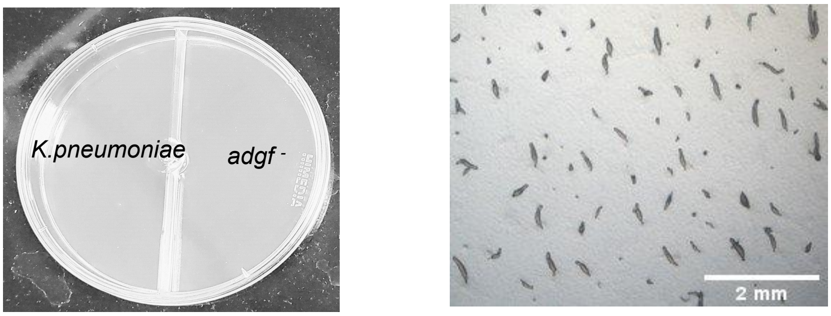


**Figure S7. Volatiles released from *K. pneumoniae* rescues the mound arrest phenotype of *adgf* -**

Picture depicting *K. pneumoniae* and *adgf* - cells on either side of a compartmentalized Petri dish. Scale bar: 2 mm; (n=3). *K. pneumoniae* and *adgf* - cells were developed on either side of compartmentalised dish, and photographs were captured at 16 h.

### Supplementary Table 1

**Various deaminases annotated in *D. discoideum***

| Gene | Gene ID |
|---|---|
| 2-aminomuconate deaminase | DDB_G0275081 |
| adenosine deaminase | DDB_G0287371 |
| adenosine deaminase acting on tRNA 1 | DDB_G0278943 |
| adenosine deaminase, tRNA-specific | DDB_G0288099 |
| adenosine deaminase-related growth factor | DDB_G0275179 |
| AMP deaminase | DDB_G0292266 |
| CMP/dCMP deaminase, zinc-binding domain-containing protein | DDB_G0282255 \| DDB_G0271914 \| DDB _G0286161 \| DDB_G0288019 |
| cytidine deaminase | DDB_G0292096 |
| cytidine deaminase-like protein | DDB_G0278841 |
| cytidine/deoxycytidylate deaminase zinc-binding domain-containing protein | DDB_G0292096 |
| dCTP deaminase | DDB_G0293580, DDB_G0268194 |
| glucosamine-6-phosphate deaminase | DDB_G0278873, DDB_G0286195 |
| guanine deaminase | DDB G0277743 |
| porphobilinogen deaminase | DDB_G0284697 |
| putative arginine deaminase | DDB_G0289195 |
| serine deaminase | DDB_G0272787 |
| threonine deaminase | DDB_G0277245 |
| glutamine synthetase | DDB_G0276835,DDB_G0295755 |
| glutamate dehydrogenase | DDB_G0280319, DDB_G0287469 |
| Glutaminase | DDB_G0291984 |
| Allantoicase | DDB_G0280267 |
| Aconitase | DDB_G0279159 |
| Arginine deiminase | DDB_G0272182 |
| Formimidoyl transferase-cyclo deaminase | DDB_G0287977 |
| Fatty acid amide hydrolase | DDB_G0275967 |

### Supplementary Table 2

Primers used for *adgf* semi-quantitative PCR

| **Primer** | **Sequence** |
|---|---|
| P1 | CCGAAGCTTAAAATGTTTTTAAAGTTTA |
| P2 | ATAGAAATGAATGGCAAGTTAG |
| P3 | GTTGAGAAATGTTAAATTGATCC |
| P4 | TGGATGAGCACGCATATCAG |

### Supplementary Table 3

Primers used for *adgf* over expression and vector construction

| **Primer** | **Sequence** |
|---|---|
| *adgf* OE FP | CCGAAGCTTAAAATGTTTTTAAAGTTTA |
| *adgf* OE RP | GGCGGTACCTTAAATATTTGAATAAGTATTA |

### Supplementary Table 4

Primers used for real time-PCR

| Primer | Sequence |
|---|---|
| *adgf* FP | GTGGTGTATGATGCAATGGTAATG |
| *adgf* RP | TGGATGAGCACGCATATCAG |
| *ada* FP | GAAACGGGTTAACTAAAGAGCAAG |
| *ada* RP | TGGATCATCAGAGGTTGAAGAAG |
| *adat* FP | ACCAATTTCAGGAGAGTGGAC |
| *adat* RP | TTCCTAAACACCTATTACCAGTACC |
| *ada*-tRNA FP | GAACAAGACACGCAGAACTTG |
| *ada*-tRNA RP | GCACATCAAACATGGCTCTAC |
| *countin* FP | CAACCGGTAATGCTTTTGGT |
| *countin* RP | CACAAACGAGAGCTGACA |
| *smlA* FP | TGGATTACACCATGTTCAGCA |
| *smlA* RP | CCGACTGAAACTGATGCTTTGG |
| *acaA* FP | CATTCTAGAGGCGGTATTGGC |
| *acaA* RP | GGAGAAAATGTCTGATTTCGCTT |
| *carA* FP | ATGTTGGGTTGTATGGCAGTG |
| *carA* RP | AGGGAAACCACCATTGACAG |
| *pdsA* FP | CCATTGGGTACAACTGGTGGA |
| *pdsA* RP | AACTGCCCATGATGGATAGGT |
| *regA* FP | TAAAGCAACGTTGGCACAAG |
| *regA* RP | ATGGTGATTCCATTGCTTCC |
| *pde4* FP | GATCTTGATACACCAATCGAA |
| *pde4* RP | CTTCTGCATCATCTGTACATG |
| *5'nt* FP | CAGCTGAACAAGTAGCAATGG |
| *5'nt* RP | TGGTGGAAGACTTGATGCTG |
| *cadA* FP | TTCCAAGAATTGGCTCAAGG |
| *cadA* RP | CATCAACTGCCCATTGAAAA |
| *csaA* FP | GCCAAATACAATCGCTGGTG |
| *csaA* RP | TGGTTGGTGTGAGATCAAAAGC |
| *ecmA* FP | CCAATTAGCTGTCCAAAACC |
| *ecmA* RP | GCAATCACCTTTACCTCCTG |
| *ecmB* FP | TGATTCATGTTGTTCAACTGG |
| *ecmB* RP | TAAATCATCGCCACATTTTCC |
| *pspA* FP | CATTGGCCAATCAAAATCCAG |
| *pspA* RP | ACAACAGTTGAAGCAGAACC |
| *rnlA* FP_qRT | TTACATTTATTAGACCCGAAACCAAGCG |
| *rnlA* RP_qRT | TTCCCTTTAGACCTATGGACCTTAGCG |
